# Cranial sexual dimorphism levels in modern humans: Supporting a global expansion originating in southern Africa

**DOI:** 10.1101/2024.02.11.579824

**Authors:** Zarus Cenac

## Abstract

Modern humans are acknowledged to have expanded across Earth from Africa. Biological measures have appeared to reflect this expansion, such as genetic diversity and cranial sexual size dimorphism. Admixture is known to be an issue for using diversity to locate where the expansion set out from in Africa; using cranial dimorphism should make admixture less of an issue. Therefore, cranial dimorphism could be of importance for clarifying the origin. This study used data sourced from the Howells dataset to infer if cranial form and shape dimorphisms indicate the expansion, and understand why cranial size dimorphism looks indicative. Form and shape dimorphisms were calculated through RMET. Size dimorphism was calculated beforehand. Cranial form and shape dimorphisms increased with distance from Africa. Form dimorphism suggested an area of origin which was predominantly in Africa and marginally in Asia. For shape dimorphism, locations spanned much farther beyond Africa. Hence, cranial form dimorphism seemed to be quite indicative of the expansion, unlike cranial shape dimorphism. Form is known to feature size and shape – cranial form dimorphism may signify the expansion mainly, or only, due to cranial size dimorphism. It seemed unsettled why cranial size and form dimorphisms seem related to the expansion because it was vague whether cranial size indicates the expansion more for males than females. Previously, a collective estimate of the origin (from biological measures including cranial size dimorphism) pointed to southern Africa; in the collective estimation process, cranial size dimorphism supported the south as would cranial form and shape dimorphisms.

## Introduction

Attempts to explain a number of occurrences in biology refer to modern humans having globally expanded from a specific continent – Africa (e.g., Ramachandran et al., 2005; von Cramon-Taubadel & Lycett, 2008). Whilst Africa does appear to be where a global expansion began (e.g., Ramachandran et al., 2005), there are different impressions of the part, or parts, of Africa in which the expansion had its origin (e.g., Manica et al., 2007; Tishkoff et al., 2009), such as the east (Ray et al., 2005) or the south (Cenac, 2023b). Nevertheless, genetic data fits with the existence of bottlenecks in the expansion (Prugnolle et al., 2005), with bottlenecks in the founding of new populations giving rise to a greatening of genetic drift (Ramachandran et al., 2005); expansion generates an explanation for why within-population diversity lowers with the build-up of geographical distance from Africa (Betti et al., 2009; Ramachandran et al., 2005), and for why the biological distance between populations greatens with the geographical distance separating populations (Ponce de León et al., 2018; Ramachandran et al., 2005) (see Figure 1).

**Figure 1.**
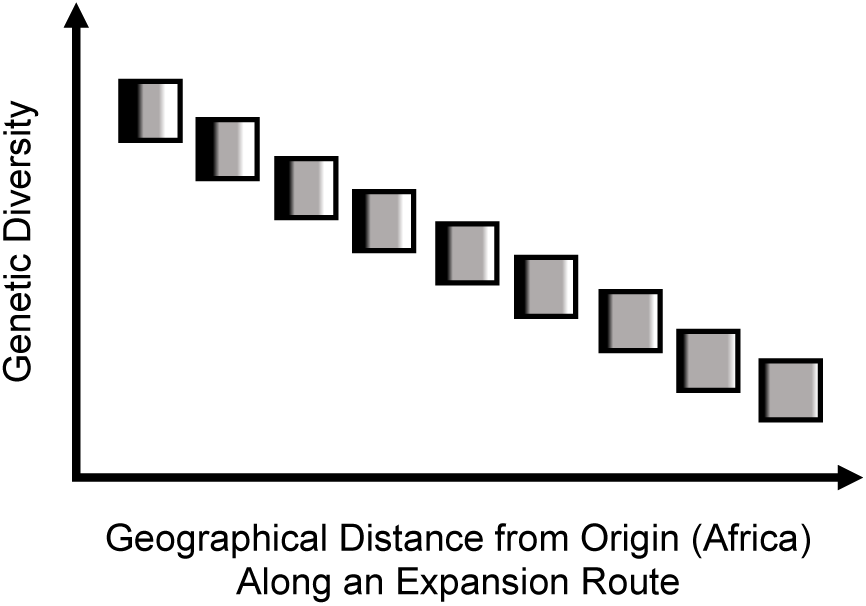
Relationship Between Genetic Diversity and Distance from Africa. *Note*. The ability of expansion to explain relationships i) between genetic diversity and how far away (geographically) populations are from Africa, and ii) between interpopulation genetic distance and geographical distance (Ramachandran et al., 2005) is represented in Figure 1. Each square is a population. In each square, diversity is embodied by the distribution of black, grey, and white. Grey gets more prevalent as geographical distance from the origin greatens (genetic drift) – genetic diversity decreases as geographical distance from Africa builds up. Research is supportive of expansion having gone along different routes (Reyes-Centeno et al., 2014), so, in Figure 1, the geographically further apart populations are in the expansion route, the greater their interpopulation genetic distance. The linear fall of genetic diversity (Prugnolle et al., 2005) is represented.

There has been discussion on what factors explain sexual dimorphism (e.g., Frayer, 1980; Smith, 1980). For instance, the popularity of sexual selection as a determiner of dimorphism has been referred to (Kleisner et al., 2021). Unpublished research has given consideration to whether sexual dimorphism is related to the expansion from Africa (Cenac, 2022a, 2022b), and such consideration was also given in the present study.^1^

### Cranial dimorphism: Size

The cranium of modern humans is sexually dimorphic in size, shape, and form (Milella et al., 2021). For cranial size dimorphism, variation between populations has not been attended to in many studies (Del Bove et al., 2023). Nevertheless, some research (unpublished) has occurred with respect to whether expansion from Africa can explain how populations vary in cranial sexual size dimorphism (Cenac, 2022a, 2022b). It has been found that cranial sexual size dimorphism has a positive correlation with geographical distance from Africa (Cenac, 2022b). Moreover, the dimorphism increases most strongly when geographical distances are measured from any one of a set of locations which cover a geographical region in Africa alone (rather than when distances are from elsewhere) (Cenac, 2022b). And so, expansion from Africa could very well have had some effect on cranial size dimorphism (Cenac, 2022a, 2022b). Clarity is wanting on why cranial size dimorphism seems attached to the expansion – it has been speculated that the relationship between distance from Africa and (adjusted for absolute latitude) cranial size dimorphism could be explained by male cranial size having changed more in the expansion than female cranial size did (Cenac, 2022a).^2^

### Cranial dimorphism: Shape and form

Previous research on the sexual dimorphism of the cranium has concerned not only cranial size, but also shape (e.g., Jantz & Ousley, 2020) and form (e.g., van Vark et al., 1989), with cranial form having both size and shape attributes (e.g., Matsumura et al., 2022). Is an increase in dimorphism with distance from Africa found when the dimorphism is of cranial shape or cranial form? Based on prior research (Cenac, 2022a, 2022b, 2023b; Hubbe et al., 2009; Kleisner et al., 2021; Matsumura et al., 2022; Messer et al., 2013), the answer might be yes.

As the geographical distance between populations rises, so too does the interpopulation distance in cranial shape (Hubbe et al., 2009). In a preprint, cranial shape distance (between populations) seemed to exhibit a stronger signal of the expansion from Africa for male crania than for female crania (Cenac, 2023b). Therefore, it should not be unforeseen for cranial shape dimorphism to be correlated with geographical distance from Africa. In terms of facial shape (which included the lower jaw, lips, etc.), previous research has found Africans (two populations) to have less sexual dimorphism than Europeans (four populations) and South Americans (two populations) (Kleisner et al., 2021); hence, given geographical distances from Africa (e.g., Li et al., 2008), one might predict a positive correlation between cranial shape dimorphism and geographical distance from Africa.

Cranial form has features of size and shape (e.g., Matsumura et al., 2022). Therefore, given that cranial size dimorphism elevates with geographical distance from Africa (Cenac, 2022b), and cranial shape dimorphism could possibly increase too, it would not be surprising if cranial form dimorphism increases with distance from Africa. Moreover, the comparatively high and low dimorphism of Pacific Islanders and Africans respectively has been noted (Messer et al., 2013) – this is consistent with cranial form dimorphism greatening in the expansion (Cenac, 2022a). Furthermore, correlation coefficients for relationships between dimorphism and distance from Africa, are, on average, positive when coefficients are averaged across cranial form dimensions (Cenac, 2022b). Consequently, cranial form dimorphism might be expected to greaten with geographical distance from Africa.

### Origin of expansion and admixture/ancestry

#### Admixture generally

Admixture is one of the issues in trying to find whichever sector of Africa the expansion set off from (Colonna et al., 2011). Diversity has been used in a number of instances to indicate the origin using different methodologies (e.g., Manica et al., 2007; Ramachandran et al., 2005; Tishkoff et al., 2009). The idea is that diversity is *greatest* where the expansion originated, and it descends gradually whilst distance from the origin increases (Ramachandran et al., 2005). So, if various locations across the world are employed like they are each the origin, then the origin of the expansion is indicated by the level of the decline in diversity (e.g., Luca et al., 2011; Ramachandran et al., 2005; Tishkoff et al., 2009) – see Figure 2; a complication with using diversity to indicate the origin is an increase in diversity because of admixture (Bergström et al., 2021). There would be no clear reason for cranial size, shape, or form dimorphisms to *generally* be increased/decreased by admixture. Therefore, cranial size dimorphism may not only be an indicator of the origin (Cenac, 2022b) but a particularly useful alternative to diversity, as may cranial shape and form dimorphisms if they too indicate the expansion.

**Figure 2.**
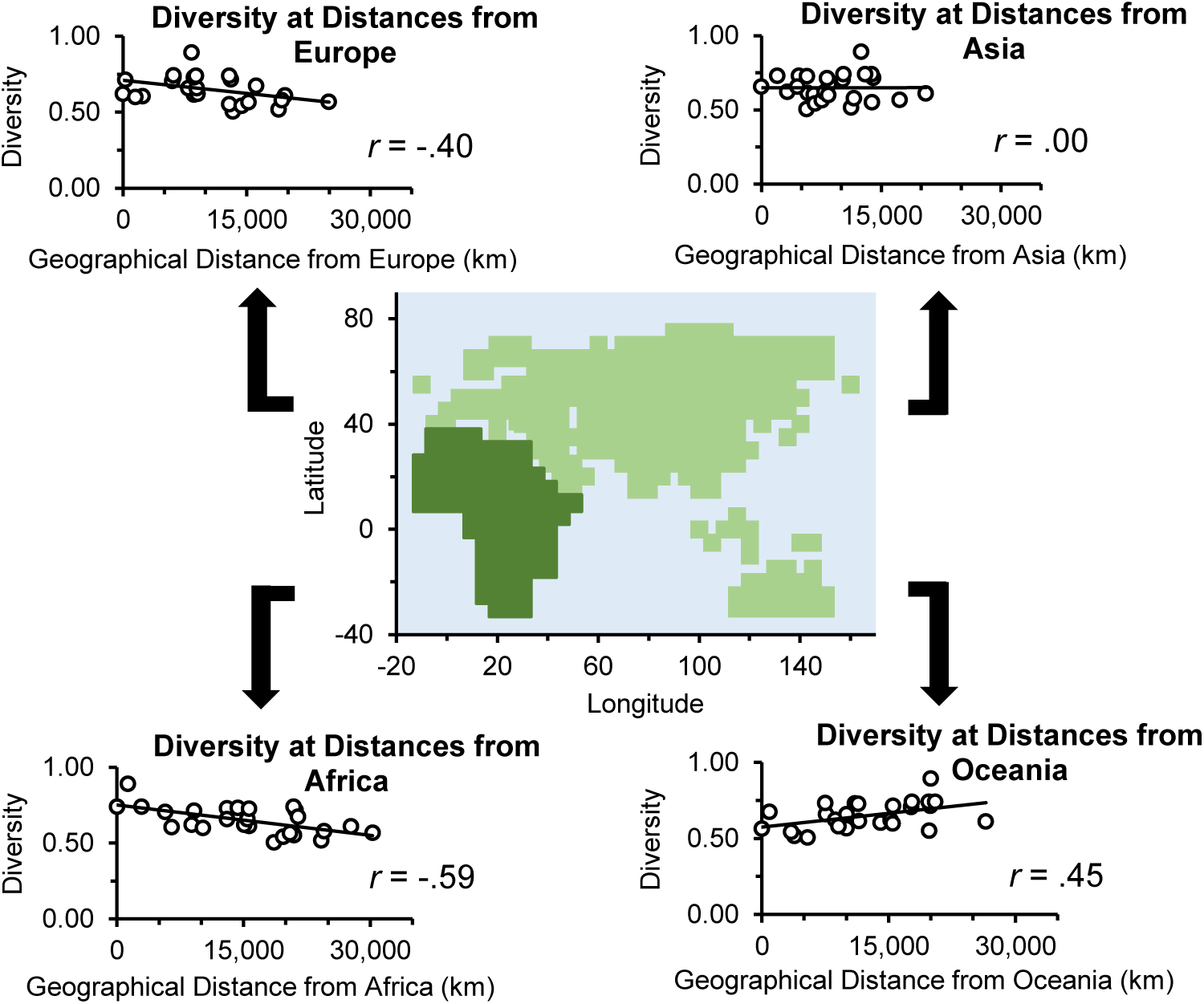
Diversity and Geographical Distance from Different Continents. *Note*. Diversity (cranial shape of females) (Cenac, 2022b) and geographical distances are from previous research (Cenac, 2022a); the Howells (1973, 1989, 1995, 1996) data were used in that research to calculate the cranial shape diversity (Cenac, 2022b), and the geographical distances are from four of the Howells data populations. The correlation coefficients had been found in Cenac (2022b), and yet they were calculated newly for Figure 2. The map uses coordinates which feature in Figure 4 of Betti et al. (2013) (the author had previously used the Africa coordinates from Betti et al., 2013, to construct figures, e.g., Cenac, 2022b). Therefore, the map is based, to some level on Betti et al. (2013) (and its colouring inspired, to an extent, on Figure 2B in Manica et al., 2007). Cranial shape diversity is known to indicate the expansion (Cenac, 2022b; von Cramon-Taubadel & Lycett, 2008). It should not be implied that the origin indicated by female cranial shape diversity spans all of mainland Africa – it does not (Cenac, 2022b).^3^

**Figure 3.**
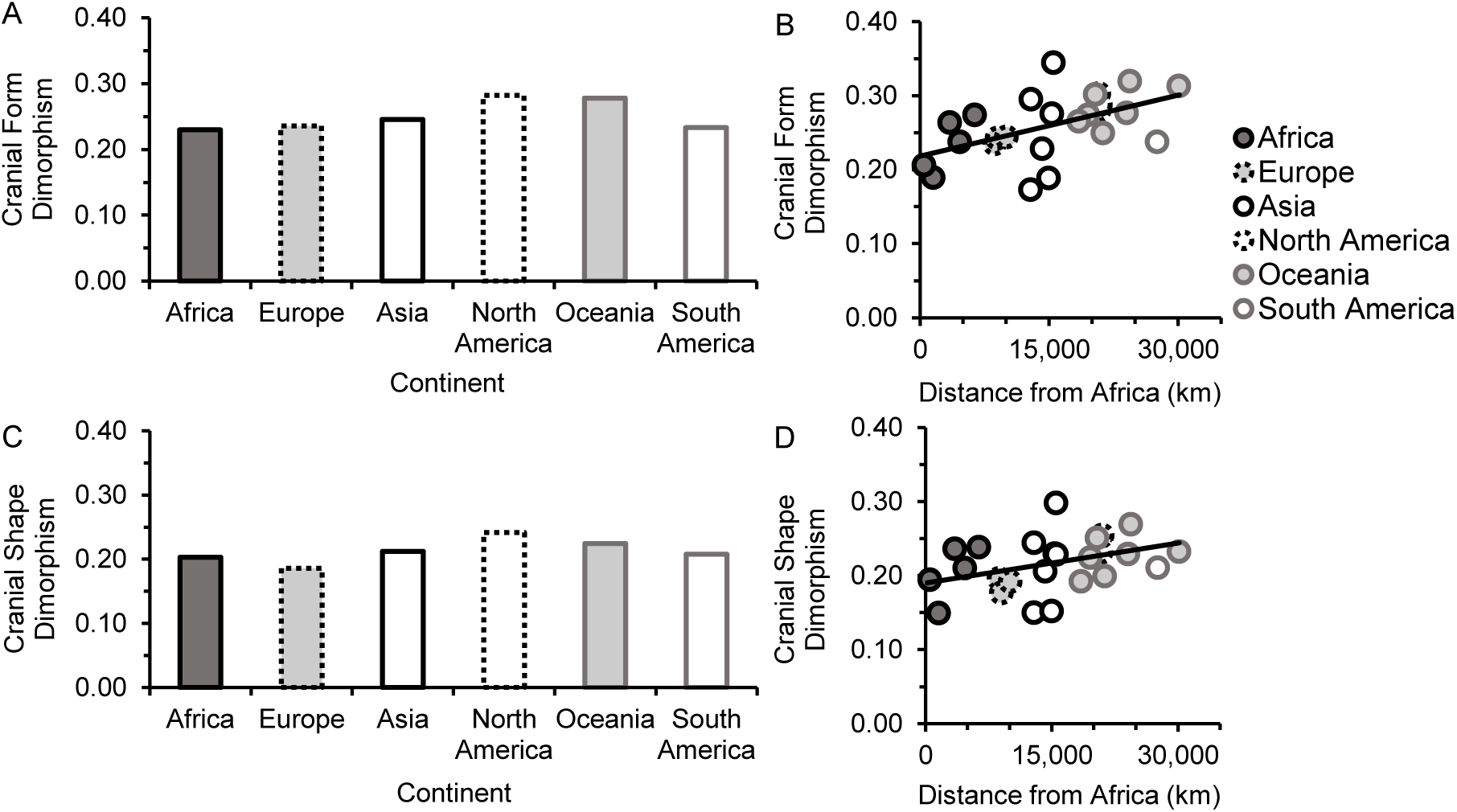
Cranial Form and Shape Dimorphisms. *Note*. Figure 3A and 3B are for cranial form dimorphism, and 3C and 3D are for cranial shape dimorphism. For 3A and 3C, dimorphisms were averaged for populations by continent. From previously (Cenac, 2022a), the continents of populations were known by the author, and the continental groupings were applied in this study. 3B and 3D show dimorphism at the population level.

**Figure 4.**
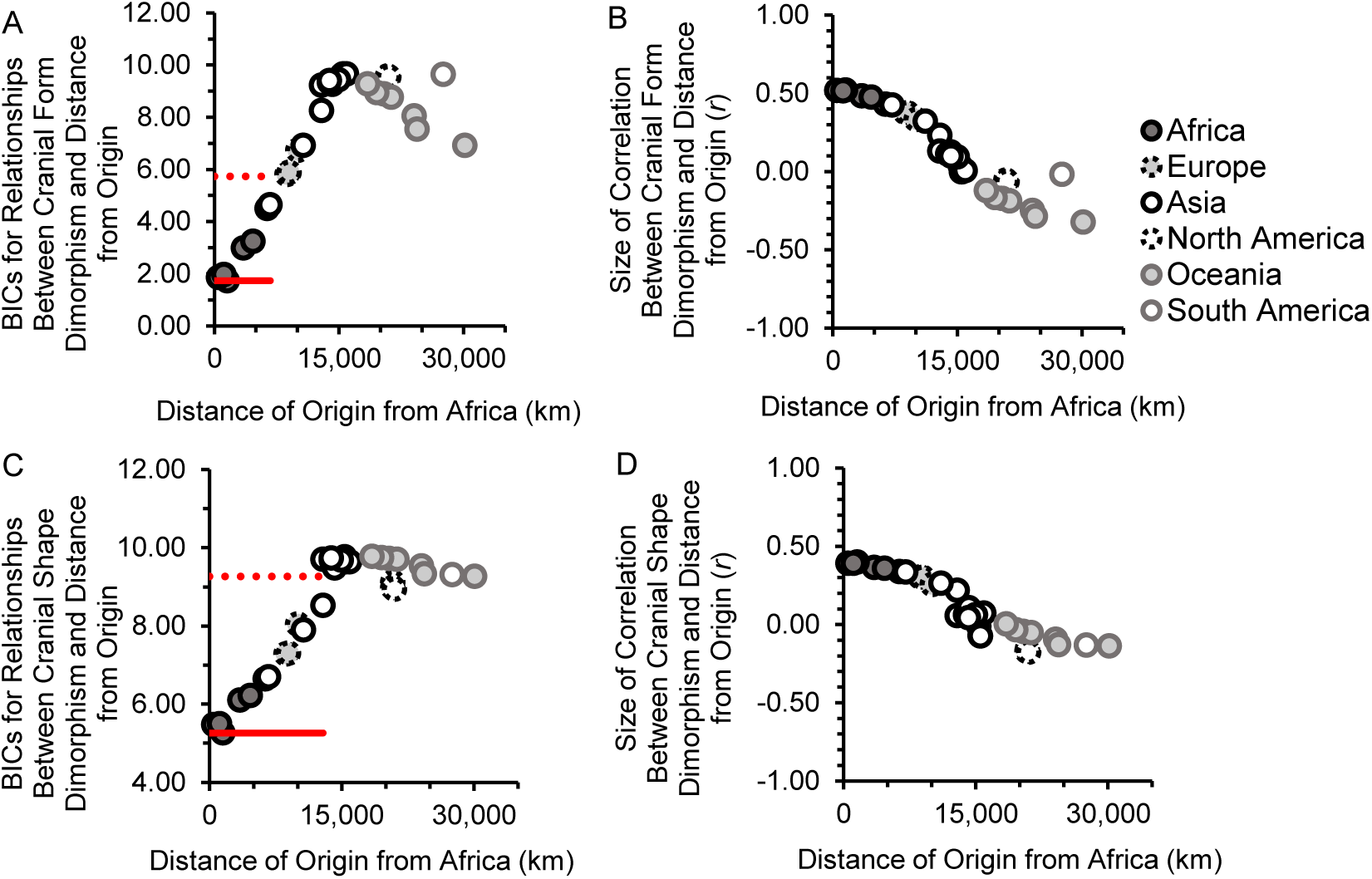
Cranial Dimorphisms in Relation to Distance from Africa. *Note*. For cranial form dimorphism (Figure 4A and 4B) and cranial shape dimorphism (4C and 4D), BICs are shown (4A and 4C) as well as correlation coefficients (4B and 4D). As before (Cenac, 2023b), the unbroken red line is the lowest BIC, whilst the dotted red line is four BICs above it – BICs from the unbroken line up and lower than the dotted line signify locations from which the increase in dimorphism is at its strongest. Figures 4 and 7 feature BICs and correlation coefficients (datapoints) for just 32 of the origins.

#### Non-African ancestry within Africa

Whilst it is acknowledged that mixing between populations usually increases diversity, results amongst Africans of admixed heritage have been in line with genetic diversity (heterozygosity) falling with rising Eurasian ancestry (Haber et al., 2016). And so, as African populations may be particular determiners of where the origin is estimated to be, it has been reasoned that non-African ancestry in Africa may have a tangible effect on the estimate – this has previously been addressed to some point by estimating the origin using autosomal diversity of only sub-Saharan Africans, with the diversity having, or not having, been adjusted for the population level of non-African ancestry (Cenac, 2022b). Cranial size dimorphism appears to track the expansion, so an estimate of the origin generated by cranial size dimorphism (Cenac, 2022b), likely would be affected by non-African ancestry in Africa. If cranial form and shape dimorphisms are linked to the expansion, then estimates of the origin derived from them would also be affected.

### This study

#### Cranial form and shape dimorphisms

A focus of the present study was on whether cranial form and cranial shape dimorphisms may be related to the global expansion. Size is a component of cranial form, as is shape (Matsumura et al., 2022), and cranial size dimorphism seems related to the expansion (Cenac, 2022b). Therefore, even if cranial form dimorphism is a product of the expansion, it would seem redundant as a measure of the expansion.

#### Cranial size

An additional focus of this study was on understanding why cranial size dimorphism is related to geographical distance from Africa (Cenac, 2022a, 2022b). As mentioned above, the relationship could be because the cranial size of females changed less in the expansion, compared to males (Cenac, 2022a). If so, this does not necessarily mean that females have less of an expansion signal in cranial size than males. For instance, even if cranial size changes more for males than females as geographical distance from Africa increases, that would not mean that a correlation between distance from Africa and cranial size is different for males than it is for females. Finding the signal to be weaker for females than males would therefore help with explaining why cranial size dimorphism correlates with distance from Africa. The idea is that cranial size dimorphism indicates the expansion (Cenac, 2022b) due to female cranial size changing less (than males) with distance from Africa (Cenac, 2022a) which reflects a weaker expansion signal in female cranial size (than males). Hence, this study examined whether an expansion signal in cranial size differs between males and females.

#### Sub-Saharan Africa and outside of Africa

To minimise a possible influence of non-African ancestry inside Africa when estimating an origin, a previous study limited some analyses solely to sub-Saharan Africans (Cenac, 2022b); in the present study, analyses regarding the area of origin were repeated (for cranial form and shape dimorphisms, and cranial size) only with populations which are either in sub-Saharan Africa or outside of Africa. Moreover, this geographical restriction was also applied when doing a re-analysis of Cenac (2022b) with respect to the origin indicated by cranial size dimorphism.

## Method

This study used cranial (form) measurements from the Howells cranial dataset (modern humans from the Holocene – 26 populations with male [1,256] and female [1,156] crania) (Howells, 1973, 1989, 1995, 1996). The dataset can be found via http://web.utk.edu/~auerbach/HOWL.htm. Additionally, cranial shape measurements of 56 dimensions from those populations in the Howells data had been calculated in prior studies (Cenac, 2022a, 2022b), and were used in the current study; the dimensions used in the present study regarding cranial form were the same 56 dimensions used concerning cranial shape. Moreover, from previous research, this study used the mean cranial sizes (mean geometric means) of males and females in the 26 populations, and cranial size dimorphism from 23 populations, and those mean sizes and size dimorphism had been calculated using measurements from the 56 dimensions in the Howells data (Cenac, 2022a). Geographical distances were sourced from preceding research (Cenac, 2022a, 2022b). All calculations occurred through R Version 4.0.5 (R Core Team, 2021), RMET 5.0 (Relethford & Blangero, 1990), and Microsoft Excel. The designs of Figures 2 to 8 are from, or based to some level on, Cenac (2022a, 2022b, 2023a, 2023b). For instance, Figures 5 and 8 have some basis in Cenac (2022b, 2023a, 2023b), who used coordinates found in a figure in Betti et al. (2013). In previous research, correlation coefficients have been represented on a map regarding i) genetic diversity and geographical distance (Luca et al., 2011; Ramachandran et al., 2005), and ii) linkage disequilibrium and geographical distance (Henn et al., 2011); the present study used such representations regarding distance from Africa and i) cranial size dimorphism (Figures 5A and 8B), ii) cranial shape dimorphism (Figure 5B), and iii) cranial form dimorphism (Figure 5C). Southern Africa was defined as a region following Choudhury et al. (2021).

**Figure 5.**
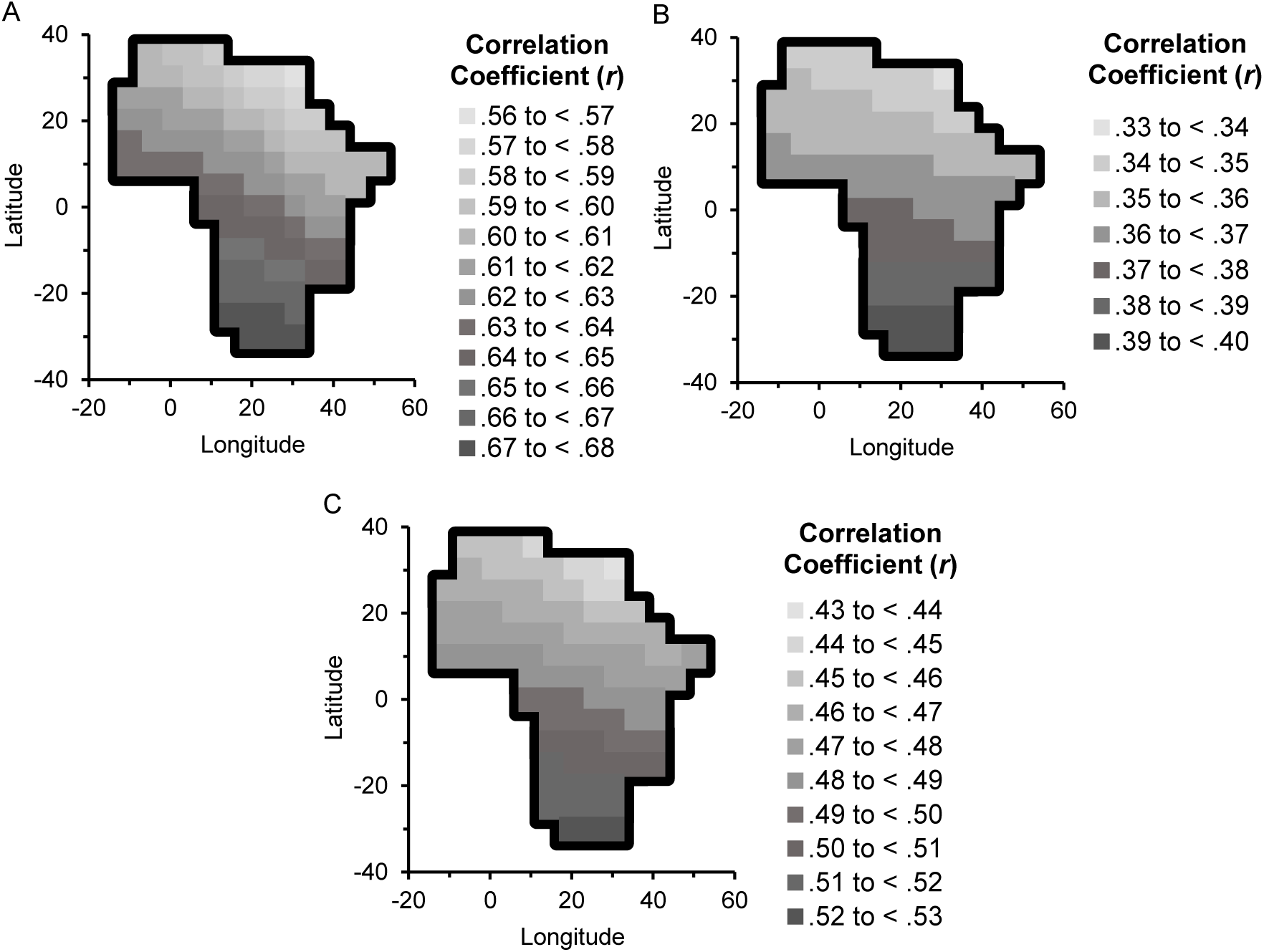
Relationships with Distance from Africa: Cranial Size, Shape, and Form Dimorphisms. *Note*. Figure 5A exhibits correlation coefficients calculated in Cenac (2023b) for relationships which are with respect to the cranial size dimorphism of 24 populations (cranial data being sourced from the Howells dataset) and geographical distance from Africa. Figure 5B has coefficients, with cranial shape dimorphism rather than cranial size dimorphism, whilst Figure 5C has coefficients regarding cranial form dimorphism and distance.

**Figure 6.**
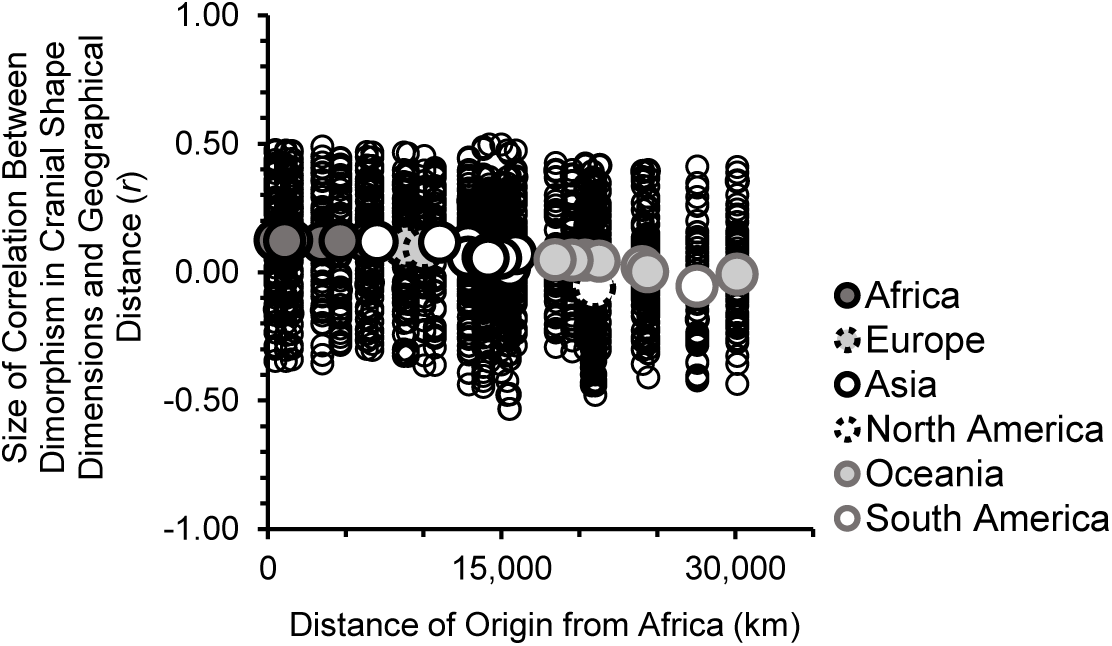
Correlation Coefficients: Cranial Shape Dimorphism of Dimensions and Geographical Distance. *Note*. For principal components derived from hip bone shape, dimorphism has been calculated as a difference between averages (males, females) (Betti, 2014); in the present study, for cranial shape dimensions, dimorphism was calculated as the absolute difference between averages – absolute values were used for consistency with overall cranial shape distances being positive distances. Smaller circles are correlation coefficients for how much the dimorphism of a shape dimension is related to geographical distance from an origin. Each larger circle is the average of correlation coefficients for an origin (i.e., across dimensions). The process of averaging correlation coefficients followed Corey et al. (1998). In Figure 6, the legend (continents) is for the larger circles.

**Figure 7.**
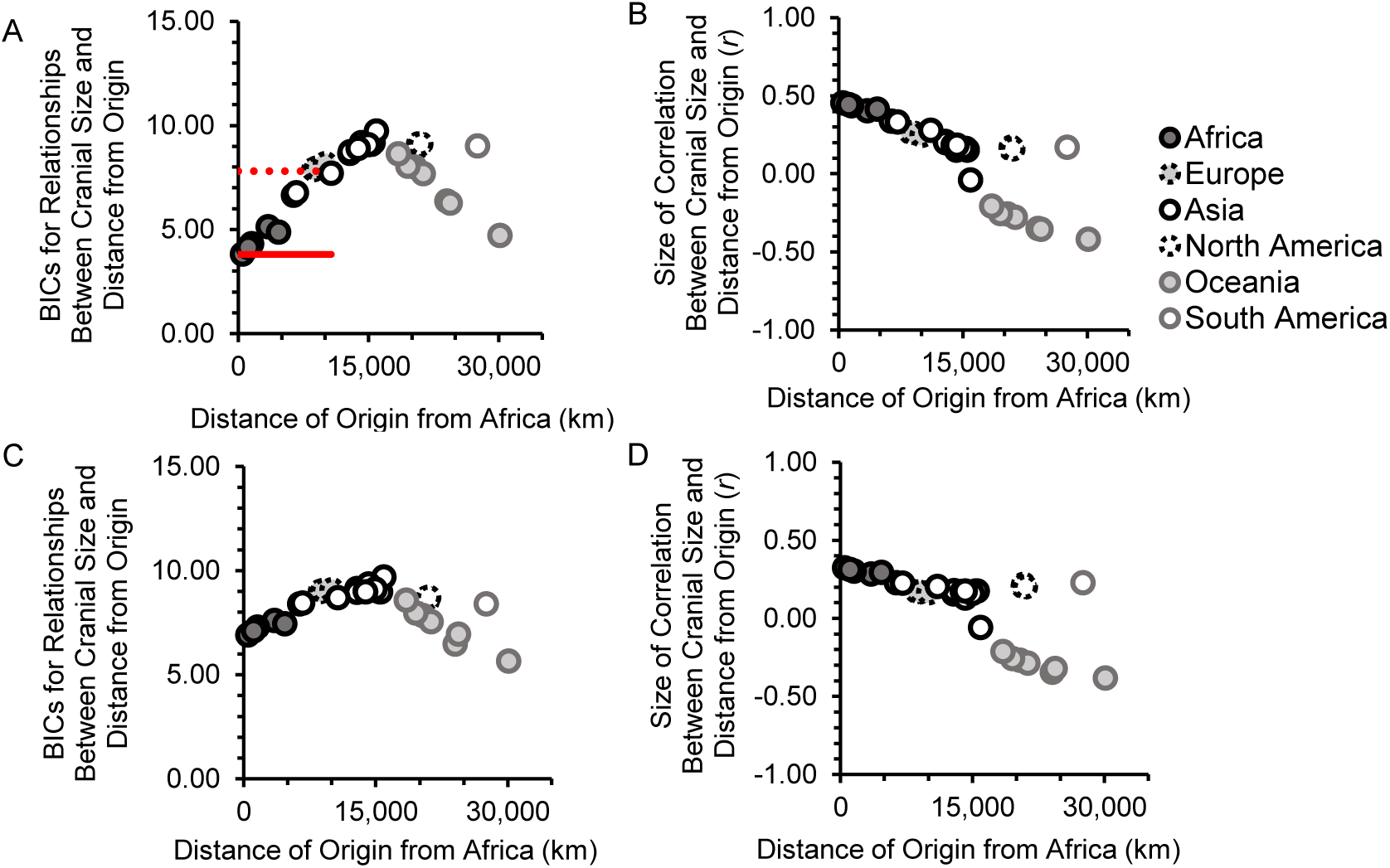
Cranial Size and Expansion. *Note*. Figure 7A and 7B are for male crania, whilst 7C and 7D are for female crania. BICs are in 7A and 7C, whilst correlation coefficients are in 7B and 7D. The red lines in Figure 7A function like the red lines in Figure 4. Here, they suggest the range of origins which gave the strongest increases in cranial size. Such lines are not shown in Figure 7C. This is because such a method of showing lines to discriminate between origins would not be clear in Figure 7C (i.e., for female crania). Indeed, BICs were without use for discriminating between origins when using female cranial size – only the sign of correlation coefficients (7D) was used.

**Figure 8.**
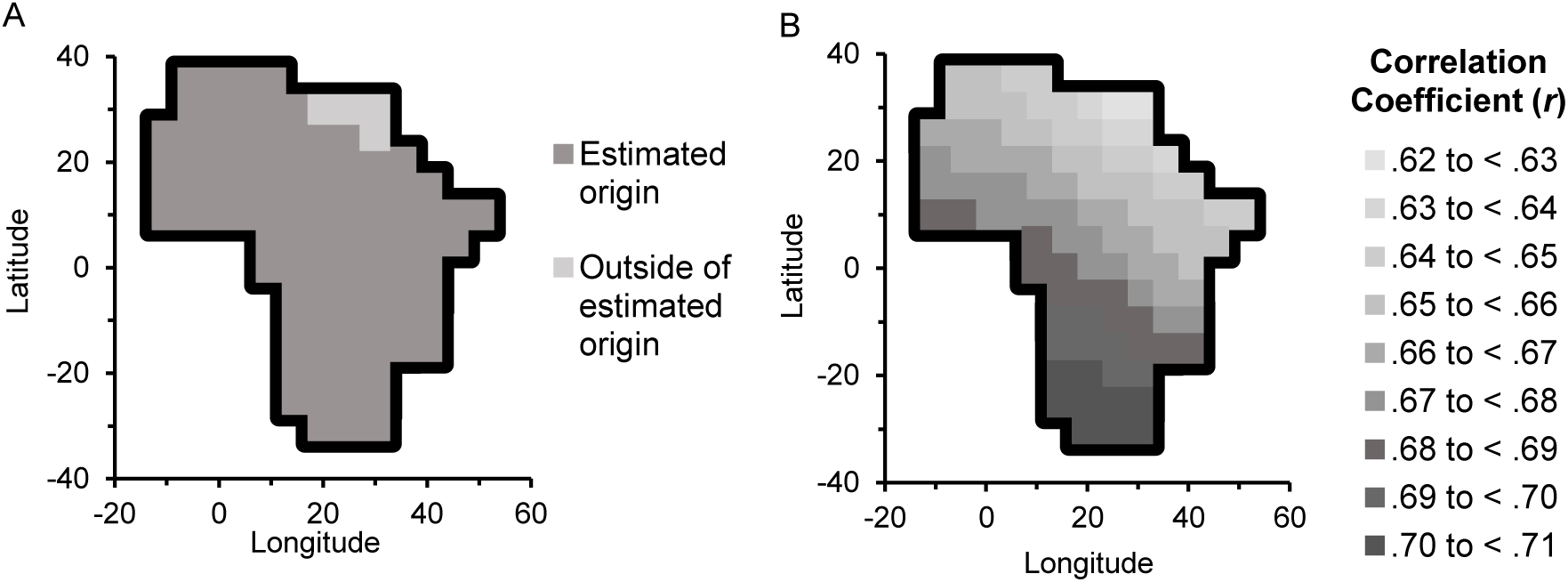
Relationships with Distance from Africa: Cranial Size Dimorphism Without Egypt. *Note*. In Figure 8A, an origin in Africa is indicated through the relationship between cranial size dimorphism and geographical distance from Africa. Figure 8B presents correlation coefficients for the relationship.

### Cranial distance

As in previous research (Pinhasi & von Cramon-Taubadel, 2009), RMET 5.0 (Relethford & Blangero, 1990) was employed to calculate squared cranial shape distances (*D*^2^). Like before (Cenac, 2023b), the square root of *D*^2^ was cranial shape distance (*D*); the measure of cranial sexual shape dimorphism was the within-population cranial shape distance between males and females. Similarly, RMET was used to calculate squared cranial form distances from which were found cranial form distances, and the cranial form distance between males and females was the measure of sexual dimorphism in cranial form. RMET had previously been used in the calculation of cranial form distances (Betti et al., 2010). As prior studies have done (Betti et al., 2010; von Cramon-Taubadel & Pinhasi, 2011), a heritability of 1.00 was used. Sexual dimorphism can be calculated in a number of ways (e.g., Messer et al., 2013; Schutz et al., 2009); the author is unaware of previous research having used *D* derived from RMET as a measure of sexual dimorphism.

### Correlation

It was tested whether geographical distance from Africa correlates with i) cranial form dimorphism, and ii) cranial shape dimorphism. Geographical distance was from the geographical location in Africa ranked highest in terms of likely origins across various cranial and genetic variables (Cenac, 2023b). Those distances were also used on the *x*-axes of Figures 3B, 3D, 4, 6, and 7. As part of the correlation analyses, it was considered whether datapoints were atypical. Atypicality was defined as a datapoint that had an absolute *z*-score standardised residual surpassing 3.29 (e.g., Field, 2013). Positive spatial autocorrelation was tested for by calculating (through a spreadsheet) a spatial Durbin-Watson (*DW*) upon which bounds for the *DW* statistic can be employed (Chen, 2016), and were indeed applied using 5% *DW* bounds from Savin and White (1977). In the event that positive spatial autocorrelation was indicated, or if it was unclear if it was indicated, the spreadsheet (including a graph) was consulted in order to suggest which population(s) the autocorrelation (or uncertainty) arose from. Because shape is in form (e.g., Matsumura et al., 2022), the correlation test regarding cranial shape dimorphism and distance from Africa was treated as a follow-up to the test concerning cranial form dimorphism and distance from Africa. Therefore, an adjustment to *p*-values (e.g., Holm, 1979) was not applied (see, e.g., Judd et al., 2009).

### Origin

The origin of the expansion has been estimated before using the Bayesian information criterion (BIC) (see Manica et al., 2007, for details). For instance, the BIC has been used regarding estimating the origin through cranial diversity (Manica et al., 2007) as well as cranial size dimorphism (Cenac, 2022b). Various locations in Africa (99 from Betti et al., 2013) and worldwide (32), including Asia (e.g., locations for Andaman, Delhi, and Tel Aviv) and Europe (von Cramon-Taubadel & Lycett, 2008), were employed like each was the geographical origin (the same origins used as in Cenac, 2022b). A BIC was found for the relationship between geographical distance from each origin, and i) cranial form dimorphism, ii) cranial shape dimorphism, iii) cranial size, and iv) cranial size dimorphism. BICs are used as a way of estimating the area in which the expansion originated from (e.g., Betti et al., 2009); the area is assembled from the origin that delivers the lowest BIC, as well as the other origins whose BICs are not dissimilar to the lowest (within four BICs) (Manica et al., 2007) – that method was used in this study. BICs were calculated via a formula (Masson, 2011).

An area of origin which is unique to the African continent (e.g., Manica et al., 2007), or predominantly there, does seem consistent with a variable being indicative of the global expansion (Betti et al., 2013). As with cranial size dimorphism beforehand (Cenac, 2022b), the present study looked for the geographical area from which there is the strongest increase in i) cranial form dimorphism, ii) cranial shape dimorphism, iii) cranial size, and iv) cranial size dimorphism – the locations giving the strongest increases would constitute the indicated area of origin; for each origin was also found the correlation coefficient for the relationship of geographical distance from the origin and each variable (cranial form dimorphism, cranial shape dimorphism, cranial size, and cranial size dimorphism).

Why look for the geographical area made of the strongest *increases*? Given reasoning in the *Introduction*, one would expect the areas of origin for cranial form and shape dimorphisms to be comprised of origins which yield positive correlation coefficients. As for cranial size, one can see a positive gradient regarding distance from Africa (*x*-axis) and cranial size (*y*-axis) (Cenac, 2022a). Therefore, as Africa is the origin of the expansion (e.g., Ramachandran et al., 2005), the area of origin made using cranial size should be composed of locations that produce positive correlation coefficients regarding cranial size and geographical distance. Cranial size dimorphism is known to rise with distance from Africa, and research has previously defined the area of origin to be made of positive correlation coefficients (Cenac, 2022b), and so the strongest increases were also looked for in the present study. Indeed, analysis with respect to cranial size dimorphism (e.g., BICs, correlation coefficients etc.) in the present study redid analysis of previous research (i.e., Cenac, 2022b), but without the Egyptian population from the Howells (e.g., 1996) data.

To infer whether the signal of the expansion is different for male cranial size than female cranial size, it was seen whether the lowest BIC for positive relationships between cranial size and geographical distance are lower for females than males or vice versa (using the four-BIC method, like what has been done with cranial shape diversity, Cenac, 2023b). Inference was also based on the indicated area of origin found through cranial size – the area found using female cranial size was descriptively compared to the area found using male cranial size.

#### Sub-Saharan Africa and outside of Africa

As noted in the *Introduction*, the process of calculating origin areas (including BICs and correlation coefficients) was, at times, done just using sub-Saharan Africans and populations beyond Africa in order for the estimate of the origin to potentially be less influenced by non-African ancestry within the African continent. In prior research, the area of origin with respect to cranial size dimorphism (and geographical distance) had been calculated in the presence of the Egyptian population (Cenac, 2022b). Origin areas were initially calculated in the present study for cranial form and shape dimorphisms, and cranial size, using the Egyptian population. However, in the current study, the area of origin (and correlation coefficients) regarding each of the cranial dimorphisms (size, form, and shape) and geographical distance, and cranial size and geographical distance, was recalculated in the absence of the Howells Egyptian population.^4^ Therefore, in Africa, only sub-Saharan African populations^5^ and populations outside of Africa (e.g., Relethford & Smith, 2018) were included in the recalculations. Relethford and Smith (2018), who also used the Howells data (but more populations in the data), also did their analyses with and without the Egyptian population.

## Results and discussion

### Cranial form dimorphism

Regarding testing whether there is a correlation between sexual dimorphism in cranial form and geographical distance from Africa, there was uncertainty over whether positive spatial autocorrelation was present, spatial *DW* = 1.44. The Ainu population (Howells, 1989, 1995, 1996) was indicated to lead to this uncertainty. Without Ainu, positive spatial autocorrelation was absent, spatial *DW* = 2.06, and there was a positive correlation between cranial form dimorphism and geographical distance from Africa, *r*(23) = .52, *p* = .008 (Figure 3B). BICs indicated that the strongest increases in cranial form dimorphism were when geographical distances were from locations in an area which was across Africa, and went into Asia partially (Tel Aviv coordinates in von Cramon-Taubadel & Lycett, 2008) (Figure 4A and 4B). So, cranial form dimorphism was not all the way congruent with indicating the global expansion, but it generally was congruent.

This can be contrasted with cranial size dimorphism, which not only increases with geographical distance from Africa, but the strongest increases in cranial size dimorphism occur only with distances from Africa (Cenac, 2022b). Therefore, cranial size dimorphism possibly has a stronger signal of the expansion (Cenac, 2022b) than cranial form dimorphism does.

### Cranial shape dimorphism

Whilst cranial shape dimorphism increased with geographical distance from Africa, *r*(24) = .39, *p* = .048, spatial *DW* = 1.78 (Figure 3D), the strongest increases were not only found when distances were from Africa, but also found for locations in Europe and some in Asia (Tel Aviv, but also Andaman and Delhi coordinates from von Cramon-Taubadel & Lycett, 2008) (Figure 4C and 4D). Therefore, it can be inferred that the area with the strongest increases was notably larger for cranial shape dimorphism than for cranial form dimorphism – it covered more of Asia, and at least some of Europe. And so, it seems particularly uncertain whether cranial shape dimorphism indicates the global expansion.

In a preprint which estimated the origin through different variables including biological diversities (Cenac, 2023b), cranial size dimorphism would have given its support to southern Africa^6^ (see Figure 5A). Cranial shape dimorphism may only be a weak indicator of the expansion, or it may not even be an indicator (Figures 3 and 4). Therefore, when using different variables to construct an estimate for the geographical source of the expansion (e.g., Cenac, 2023b), it would seem pointless to include cranial shape dimorphism. Nevertheless, for cranial shape dimorphism, southern Africa appears to be where the numerically highest correlation coefficients are (Figure 5B). Therefore, inclusion of cranial shape dimorphism in the cross-variable estimate would give further favour to the south being the origin. Similarly, if a cross-variable estimate (e.g., Cenac, 2023b) included cranial form dimorphism, then cranial form dimorphism would seem to complement support for southern Africa. However, cranial form is comprised of not only size but also shape (Matsumura et al., 2022), meaning that cranial form dimorphism would be redundant to cranial size and shape dimorphisms when estimating the origin. Given a general indication of a southern origin (Cenac, 2023b), finding the numerically highest coefficients to be in the south for cranial shape dimorphism does add some credence to the expansion being represented, to some level, in cranial shape dimorphism.

In the context of a prior study (Milella et al., 2021), perhaps it should not be surprising for there to be a weak, or absence of, indication of the global expansion in cranial shape dimorphism. In that study, which used crania of Italians, there was a significant difference between males and females in cranial form, size, and shape (Milella et al., 2021). However, whilst sex explained 18% of the variance in cranial form, and 46% of the variance in cranial size, it explained just 2% of the variance in cranial shape (Milella et al., 2021). Indeed, there was a notable coinciding in cranial shape between males and females (Milella et al., 2021). Therefore, if cranial shape dimorphism truly does hold a signal of the expansion from Africa, it should not be unforeseen for this signal to be weak.

So, which type of cranial dimorphism (size, shape, or both) is responsible for cranial form dimorphism appearing to be quite in sync with the expansion from Africa? Cranial size dimorphism and the expansion seem to be associated (Cenac, 2022b), and there is a vagueness regarding the situation with cranial shape dimorphism (Figure 4C and 4D), meaning that the answer likely is: cranial size dimorphism, either principally or only.

Across cranial dimensions, the correlation between cranial form dimorphism and geographical distance does appear to generally have its *numerical* strongpoint when Africa is where geographical distances are from (Cenac, 2022b; see also Cenac, 2022a). This also happens regarding the dimorphism of cranial shape dimensions (Figure 6). Compared to correlation coefficients found with cranial dimensions for form dimorphism and distance from Africa (Cenac, 2022b), fewer coefficients would seem to be positive for relationships between the shape dimorphism of dimensions and distance from Africa (Figure 6); perhaps the correlation between cranial shape dimorphism and distance from Africa mainly results from just a few shape dimensions.

### Cranial size

Could the signal of expansion from Africa be stronger in cranial size for males than females? Regarding cranial size and geographical distance from origins, for origins delivering positive correlation coefficients, the lowest BIC for males was in Africa, as was the lowest BIC for females. Overall, for origins giving positive correlation coefficients, the lowest of the BICs was for males, however, the lowest BIC for females was only 3.13 BICs above it. Therefore, if there is an expansion signal in cranial size, BICs suggested that the signal is not substantially greater for males than females. BICs, and also correlation coefficients, were used to indicate the geographical area of origin (Figure 7). For males, the area was in Africa and Asia – all of the African origins were in the area, along with two origins outside of Africa, which were Delhi and Tel Aviv. For females, BICs had no value in distinguishing an area of origin, leaving only correlation coefficients as the determiner – the area (positive coefficients) was across Africa, in Asia, Europe, North America, and South America. This is suggestive of a weaker expansion signal in cranial size for females than males, or even an absence of an expansion signal in female cranial size.

These analyses regarding cranial size were done with 26 populations. However, the correlation of cranial size dimorphism and distance from Africa was found using 24 populations, with two populations omitted due to positive spatial autocorrelation (Cenac, 2022b). In the present study, when the analyses regarding cranial size were repeated using the 24 populations, results were similar to when 26 were used.

All in all, a comparison of the lowest BICs for males and females is not supportive of the expansion signal being greater for males, whereas descriptive analysis with the area of origin does appear supportive. Consequently, it seems unclear whether an expansion signal in cranial size (if there is one) can be characterised as being stronger for the crania of males than the crania of females.

### Sub-Saharan Africa and outside of Africa

The areas of origin found with dimorphism in cranial size (Cenac, 2022b), form, and shape (present study), and cranial size, were recalculated in the absence of the Egyptian population in order to reduce the possible impact of non-African ancestry with respect to origin estimation (see *Introduction* and *Method*). Without the Egyptian population, the areas of origin suggested by cranial form dimorphism and cranial shape dimorphism were indicated to match the areas found when the Egyptian population was included. As for cranial size, areas of origin were generally similar to those found with the inclusion of the Egyptian population. However, the area found using male crania of the now 25 populations was smaller than when 26 populations were used, with Delhi no longer being in the area of origin.

Previously, when including the Egyptian population, cranial size dimorphism has produced an area of origin which was not anywhere apart from Africa (Cenac, 2022b). Without the Egyptian population, it can be seen that the area of origin is still completely within Africa (Figure 8A). Moreover, even without that population, correlation coefficients still indicate that cranial size dimorphism would be supportive of southern Africa being the origin (Figure 8B) if cranial size dimorphism is featured when constructing a collective estimate of the origin with biological variables like in Cenac (2023b); this support for a southern origin is likely not because of non-African ancestry. This should add some further confidence to the south of Africa being the geographical region which the global expansion set out from.

### Speculations and future research

#### Cranial shape dimorphism: Accounting for diversity

Cranial shape diversity appears to signify expansion from Africa, declining as geographical distance from the continent increases whether for males (von Cramon-Taubadel & Lycett, 2008) or females (Cenac, 2022b). In the present study, the measure of cranial shape dimorphism did not take into account the decline of diversity. It is possible to take diversity into account when calculating dimorphism (Schutz et al., 2009). A method which does that (Schutz et al., 2009), and has been used amongst humans regarding hip bone shape dimorphism, calculates dimorphism by finding a squared Procrustes distance (between males and females) and dividing it by the sum of variances for males and females (Betti, 2014). Squared Procrustes distances were not used in the present study. Nevertheless, in future research, it may be desirable to use such a method (dividing distance by summed diversity) (Schutz et al., 2009) when calculating cranial shape dimorphism. Given the decline in cranial shape diversity (Cenac, 2022b; von Cramon-Taubadel & Lycett, 2008) and the increase in cranial shape dimorphism with distance from Africa (Figure 3D), it could be argued that results in the present study are likely to have underestimated the expansion signal in cranial shape dimorphism.

#### Selection

To reiterate, the expansion appears to be represented less for females than males in cranial shape (Cenac, 2023b), and it is not clear if this can also be said for cranial size (present study). Extending some findings and discussion from Zichello et al. (2018), perhaps future research might consider whether selection in the global expansion could have resulted in the expansion being represented more strongly in male crania than in female crania.

In Zichello et al. (2018), for hominoids, genetic variation between taxa seemed to be reflected less readily in cranial shape for females than males. This could possibly be due to selection having acted to hamper cranial shape variation of females in some taxa (compared to males) (Zichello et al., 2018). Could this *selection* theory of Zichello et al. (2018) be applied amongst modern humans with respect to cranial shape and cranial size? If yes, then, compared to male crania, female crania would be i) less diverse and ii) show genetic distances less well.

Amongst *Homo sapiens*, cranial shape diversity does not appear to be smaller for females than males (Zichello et al., 2018). Indeed, for modern humans, cranial shape diversity is no different for females than it is for males (Milella et al., 2021). What about genetic distances? There is a correlation between interpopulation distance in cranial shape and genetic distance (Harvati & Weaver, 2006). Geographical distance has been utilised in place of genetic (Hubbe et al., 2009); interpopulation cranial shape distance and interpopulation geographical distance are correlated (Hubbe et al., 2009) and not more weakly for females compared to males (Cenac, 2023b). This, therefore, indicates that the correlation between cranial shape and genetic distances (Harvati & Weaver, 2006) would not be lower for females than males. Hence, it seems unlikely that the logic of Zichello et al. (2018) can be extended to explain why previous research (Cenac, 2023b) found an indication that a weaker global expansion signal may be present in cranial shape for females than males.

Can the Zichello et al. logic be applied to cranial size? The cranial size diversity of modern humans is greater for males than females (Milella et al., 2021). However, a relationship would not seem to be present between interpopulation cranial size distance and genetic distance (Harvati & Weaver, 2006). Nevertheless, the expansion from Africa could still have resulted in more of a change for males than females in cranial size (Cenac, 2022a), which, assuming that geographical distance can be used to represent genetic distance (Hubbe et al., 2009), may i) suggest a greater change in cranial size for males than females as genetic distance from Africa increases, and ii) therefore indicate that genetic distance (from Africa) is shown in cranial size at least for males. Nonetheless, it appears uncertain if cranial size has a greater global expansion signal amongst males than females (present study). What if the signal is stronger for males? If so, then, extending the reasoning from Zichello et al. (2018), research could focus on whether selection is responsible.

### Conclusion

For modern humans, there does appear to be a pattern to interpopulation variation in cranial dimorphism for size (Cenac, 2022b), form, and shape (Figure 3). For populations, their geographical distance from Africa has relationships with cranial size dimorphism (Cenac, 2022b), cranial form dimorphism, and cranial shape dimorphism (Figure 3). Nonetheless, the global expansion from Africa seems to be represented in cranial size dimorphism (Cenac, 2022b), but perhaps less so in cranial form dimorphism, and even less (or not represented) in cranial shape dimorphism (Figures 3 and 4); if cranial form dimorphism is connected to the global expansion from Africa, then this connection may be more, or solely, due to cranial size dimorphism rather than cranial shape dimorphism. However, the reason for why there is a possible expansion signal in cranial size dimorphism (Cenac, 2022b), and therefore cranial form dimorphism, is unclear given that it is uncertain if the expansion signal is greater in cranial size for males than females. The origin of expansion indicated by cranial dimorphism should be less affected by admixture than the origin indicated by diversity (see *Introduction*). Therefore, cranial size dimorphism may be particularly useful for deducing the segment of Africa in which began the global expansion. Indeed, a cross-variable estimate from biological variables backs the south (Cenac, 2023b), and cranial size dimorphism (in the estimate) is inclined to the south (e.g., Figure 5A), which supports the cross-variable estimate.

## Footnotes

1 Parts of the current study were presented in a poster by the author at the Anatomical Society Winter Meeting of 2024 (Cenac, 2024).

2 A correlation has not been evident between geographical distance from Africa and (adjusted for absolute latitude) cranial size (Cenac, 2022a). This does likely suggest that there is no correlation between (unadjusted) cranial size and distance from Africa. Testing for such a correlation was not of interest in the present study. Nonetheless, the expansion may be expressed better through tests of whether distance from Africa correlates with cranial size dimorphism than through tests of whether that distance correlates with cranial size (with size dimorphism and size adjusted for absolute latitude) (Cenac, 2022a). It has been implied/speculated that this could be because a potential relationship between climate and size may be better accounted for by using an (adjusted) cranial size dimorphism variable than an (adjusted) cranial size variable (Cenac, 2022a). The correlation of adjusted dimorphism and distance from Africa could indicate a greater shift in size for males than females in the expansion (Cenac, 2022a). If there is a greater change for males (Cenac, 2022a), this suggests that the cranial size of males does actually change with geographical distance from Africa. And so, analysis with cranial size dimorphism (Cenac, 2022b) may suggest an expansion signal in cranial size for males at least.

3 Von Cramon-Taubadel and Lycett (2008) presented cranial shape diversity of males at geographical distances from Africa and Asia.

4 Research points to the crania of the Howells Egyptian population having a kinship with populations beyond the African continent (Algee-Hewitt, 2011).

5 The Howells dataset features Dogon, Teita, San, and Zulu populations (Howells, 1989, 1996). These are sub-Saharan African populations (e.g., Relethford & Smith, 2018). When adding up ancestries presented in Tishkoff et al. (2009), about half of the ancestry of Dogon appears to be non-African (Cenac, 2022b). However, Tishkoff et al. used just nine Dogon. Across 24 Dogon, however, African ancestry is indicated to be between 98% and 100% (Xing et al., 2010). Teita are with respect to Kenya (e.g., Relethford, 2004). Kenyan Africans have predominantly African ancestry (Tishkoff et al., 2009). The ancestry of San is largely African (Tishkoff et al., 2009), as is the ancestry of Zulu at the least (Gurdasani et al., 2015).

6 The highest correlation coefficient for the relationship of cranial size dimorphism and distance is when distance is from coordinates in southern Africa (Cenac, 2022b). There does seem to be a geographical pattern to how good geographical locations are at being candidates for the origin (e.g., Ramachandran et al., 2005). Therefore, it seems likely that cranial size dimorphism favoured southern Africa in the estimation.

